# Reduced Liver-Specific PGC1a Increases Susceptibility for Short-Term Diet-induced Weight Gain in Male Mice

**DOI:** 10.1101/2020.12.04.412460

**Authors:** E. Matthew Morris, Roberto D. Noland, Michael E. Ponte, Michelle L. Montonye, Julie A. Christianson, John A. Stanford, John M. Miles, Matthew R. Hayes, John P. Thyfault

## Abstract

Central integration of peripheral neural signals is one mechanism by which systemic energy homeostasis is regulated. Previous work described increased acute food intake following chemical reduction of hepatic fatty acid oxidation and ATP levels, which was prevented by common hepatic branch vagotomy (HBV). However, possible offsite actions of the chemical compounds confound the precise role of liver energy metabolism. Herein, we used a liver-specific PGC1a heterozygous (LPGC1a) mouse model, with associated reductions in mitochondrial fatty acid oxidation and respiratory capacity, to assess the role of liver energy metabolism in systemic energy homeostasis. LPGC1a male mice have 70% greater high-fat/high-sucrose (HFHS) diet-induced weight gain and 35% greater positive energy balance compared to wildtype (WT) (p<0.05). The greater energy balance was associated with altered feeding behavior and lower activity energy expenditure during HFHS in LPGC1a males. Importantly, no differences in HFHS-induced weight gain or energy metabolism was observed between female WT and LPGC1a mice. WT and LPGC1a mice underwent sham or HBV to assess whether vagal signaling was involved in HFHS-induced weight gain of male LPGC1a mice. HBV increased HFHS-induced weight gain (85%, p<0.05) in male WT, but not LPGC1a mice. As above, sham LPGC1a males gain 70% more weight during short-term HFHS feeding than sham WT (p<0.05). These data demonstrate a sexspecific role of reduced liver energy metabolism in acute diet-induced weight gain, and the need of more nuanced assessment of the role of vagal signaling in short-term diet-induced weight gain.

**Key Points Summary:**

- Reduced liver PGC1a expression results in reduced mitochondrial fatty acid oxidation and respiratory capacity in male mice.
- Male mice with reduced liver PGC1a expression (LPGC1a) demonstrate greater short-term high-fat/high-sucrose diet-induced weight gain compared to wildtype.
- Greater positive energy balance during HFHS feeding in male LPGC1a mice is associated with altered food intake patterns and reduced activity energy expenditure.
- Female LPGC1a mice do not have differences in short-term HFHS-induced body weight gain or energy metabolism compared to wildtype.
- Disruption of vagal signaling through common hepatic branch vagotomy increases short-term HFHS-induced weight gain in male wildtype mice, but does not alter male LPGC1a weight gain.

## Introduction

Understanding the homeostatic mechanisms that mediate susceptibility to weight gain is becoming more critical as the prevalence of obesity continues to rise (Ogden *et al*., 2020). Indeed, prevention of weight gain may be a more successful strategy to combat the obesity epidemic in light of the evidence that losing and maintaining body weight is difficult (Hill *et al*., 2012). Considering the etiology of overweight/obesity, it is worth noting that long-term weight gain commonly results as the culmination of small, sporadic bouts of weight gain occurring over long time frames (Yanovski *et al*., 2000; Hull *et al*., 2006; Ma *et al*., 2006; Racette *et al*., 2008) as a function of metabolic, hedonic, hormonal, and satiety driven mechanisms (Saper *et al*., 2002; Galgani & Ravussin, 2008; Hariri & Thibault, 2010; Hopkins *et al*., 2011). Among these mechanisms is a growing list of peripheral hormonal and afferent neural signals that are integrated in the brain and impact whole-body energy homeostasis (Kim *et al*., 2018).

The regulation of energy balance is complex and multifactorial, with energy intake and energy expenditure being independently and dependently regulated (Hand *et al*., 2015). Both energy intake and expenditure are modulated by the central action of peripheral hormones (e.g. – leptin, ghrelin, & insulin), and integration of peripheral neural signals. Previous findings showed that inhibition of hepatic fatty acid oxidation with intraperitoneally injected chemical inhibitors acutely increased food intake (Scharrer & Langhans, 1986; Langhans & Scharrer, 1987; Friedman *et al*., 1999; Horn *et al*., 2001; Horn *et al*., 2004). Further, these findings suggested that liver energy metabolism could influence energy intake via afferent vagal signals (Friedman & Sawchenko, 1984; Langhans & Scharrer, 1987; Friedman *et al*., 1999; Horn *et al*., 2004), as the hyperphagic effects produced by reduced hepatic fatty acid oxidation could be eliminated if vagal communication was disrupted by common hepatic branch vagotomy (Langhans & Scharrer, 1987; Horn *et al*., 2001). However, these findings were potentially confounded by off-target effects of the chemical inhibitors.

To more specifically target liver energy metabolism with molecular techniques, we utilized a mouse model with a liver-specific reduction in the *pgc1a* gene (LPGC1a) (Estall *et al*., 2009). PGC1a is a key regulator of mitochondrial biogenesis and fatty acid oxidation in the liver (Morris *et al*., 2012) and serves as a transcriptional node for energy/nutrient signaling pathways such as fasting and reduced energy status (ATP/ADP ratio) (Handschin & Spiegelman, 2006; Fernandez-Marcos & Auwerx, 2011).The LPGC1a mouse has reduced expression of hepatic fatty acid oxidation genes (Estall *et al*., 2009), opposite the effect we have previously reported with hepatic PGC1a overexpression which increased fat oxidation and mitochondrial capacity (Morris *et al*., 2012). Using the LPGC1a mouse we tested the hypothesis that reduced liver mitochondrial fatty acid oxidation and respiratory capacity would result in greater weight gain during exposure to short-term high-fat, high-sucrose feeding in male and female mice. Our findings support an association of decreased liver energy metabolism and dysregulation of energy homeostasis during short-term high-fat, high-sucrose feeding in male, but not female, mice. Also, additional common hepatic branch vagotomy experiments support the role of vagal nerve in energy homeostasis, but do not clarify whether communication of liver energy metabolism to the brain is a necessary component.

## Methods

### Animals

The animal protocol was approved by the Institutional Animal Care and Use Committee at the University of Missouri, Harry S Truman Memorial Veterans’ Hospital, and University of Kansas Medical Center. All experiments were carried out in accordance with the *Guide for the Care and Use of Laboratory Animals* published by the US National Institutes of Health (NIH guide, 8th edn, 2011). Mice were anaesthetized with pentobarbital sodium (75 mg/kg) before a terminal procedure. Mice with liver-specific, PGC1a heterozygosity (LPGC1a) were produced as previously described (Estall *et al*., 2009; Fletcher *et al*., 2018; Von Schulze *et al*., 2018). Briefly, C57Bl/6J male mice (#000664, Jackson Laboratory, Bar Harbor, ME, USA) were bred to female homozygous *pgc1a* floxed mice (#00966, B6N.129(FVB)-*Ppargc1a^tm2.1Brsp^*/J, Jackson Laboratory, Bar Harbor, ME, USA) to produce heterozygous *pgc1a* floxed offspring. Female heterozygous *pgc1a* floxed mice were subsequently bred to male mice with transgenic expression of *cre* recombinase gene under control of the albumin promoter (#003574, B6.Cg-*Speer6-ps1^TG(Alb-cre)21Mgn^*/J, Jackson Laboratory, Bar Harbor, ME, USA). The resultant littermates all express the albumin-cre transgene, and are either homozygous (wildtype, +/+) or heterozygous for *pgc1a* in a liver-specific manner (LPGC1a, +/-). Mice were housed at ~25-27°C on 12/12 light/dark cycle, with *ad lib* access to water and low-fat diet (LFD; D12110704: 10% kcal fat, 3.5% kcal sucrose and 3.85 kcal/g energy density, Research Diets, New Brunswick, NJ, USA). All mice were fasted for 4 hr prior to euthanasia and tissue collection. Livers were quickly removed following exsanguination, and either stored in ice cold homogenization buffer (100 mM KCl, 40 mM Tris·HCl, 10 mM Tris-base, 5 mM MgCl_2_·6H_2_O, 1 mM EDTA, and 1 mM ATP; pH 7.4) or snap frozen in liquid N_2_ for storage at −80°C.

### Genotyping & RT-PCR

Mouse and liver-specific genotypes were confirmed as previously described (Lin *et al*., 2004; Estall *et al*., 2009). A list of appropriate genotyping primers is provided in Table 1, and representative images of the genotyping are presented in Figure 1A. RNA was isolated from ~20-30 mg of liver tissue using the RNeasy Plus Mini Kit (Qiagen, Valencia, CA, USA), with cDNA produced using the ImProm-II RT system (Promega, Madison, WI, USA). Real-time quantitative PCR was performed using a Prism 7000 (Applied Biosystems, Foster City, CA, USA) and SYBR Green. The SYBR Green primers for PGC1a are provided in Table 1. All gene specific values are normalized to Cyclophilin B (*ppib*) expression.

**Figure 1.**
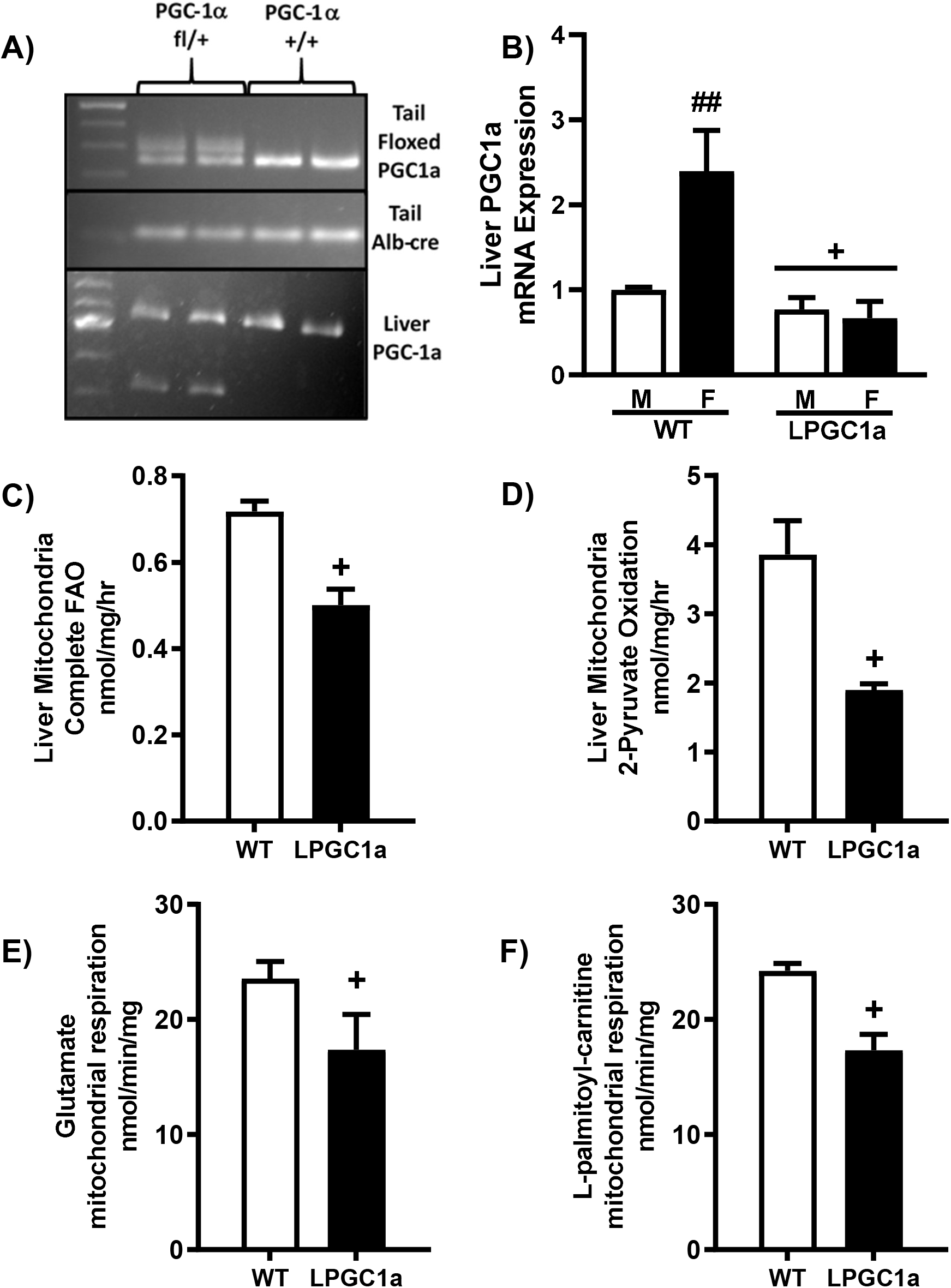
Liver-specific PGC1a heterozygosity results in reduced fatty acid oxidation and oxidative capacity in isolated liver mitochondria. A) Tail and liver genotyping of WT and LPCG1a mice. B) Relative liver mRNA expression of PGC1a. Complete oxidation of 1-[14C]-palmitate (C) and 2-[14C]-pyruvate (D) to CO_2_ in isolated liver mitochondria in male mice. Isolated liver mitochondrial respiratory capacity was determined in male mice by measurement of O_2_ consumption using a Clark electrode system in the presence of (E) glutamate (+malate) and (F) L-palmitoyl-carnitine (+malate) during state 3 respiration (+ADP). Values are means ± SEM (n= 4 - 6). + p<0.05 WT vs. LPGC1a. ## p<0.05 male vs. female within genotype.

**Table 1.**
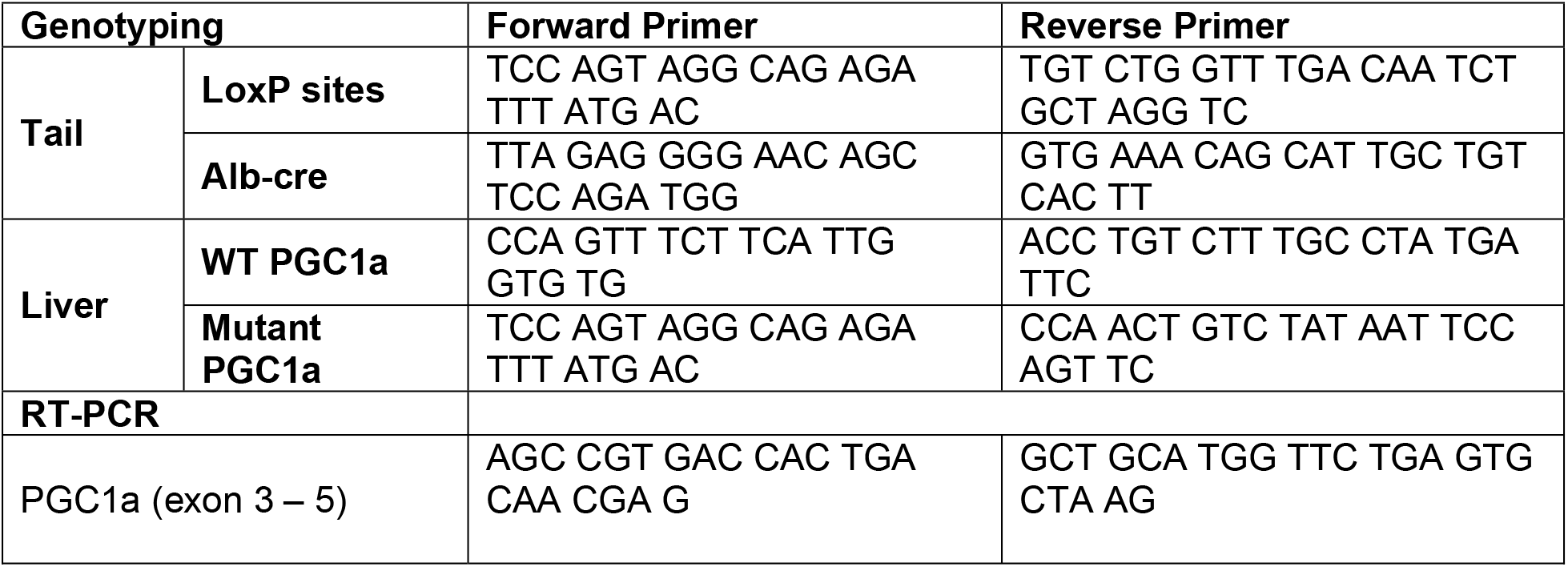
Primers

### Fresh Liver Mitochondria Isolation

Isolated liver mitochondrial fatty acid and pyruvate oxidation was performed as previously described (Morris *et al*., 2012). To isolate liver mitochondria, approximately 1 g of liver was crude minced in 8 mL of ice-cold mitochondrial isolation buffer (220 mM mannitol, 70 mM sucrose, 10 mM Tris, 1 mM EDTA, pH adjusted to 7.4 with KOH). Minced tissue was homogenized with a 15 mL glass-on-teflon homogenizer (8 passes @ 2,000 rpm). Crude homogenate was created by centrifugation (4°C, 10 min, 1500 g). The supernatant was transferred to a round bottom tube and centrifuged (4°C, 10 min, 8000 x g). The pellet was resuspended in 6 mL of isolation buffer using a Dounce glass-on-glass homogenizer and centrifuged again (4°C, 10 min, 6000 x g). The pellet was resuspended in 4 mL of isolation buffer containing 0.1% BSA and centrifuged (4°C, 10 min, 4000 x g). This final pellet was resuspended in ~0.75 mL of isolation buffer. The protein concentration for both suspensions was determined by BCA assay.

### Fatty Acid and Pyruvate Oxidation

Complete oxidation of [1-14C]-palmitate and [2-14C]-pyruvate to CO_2_ was measured in liver mitochondria (n=4, male), as previously described (Morris *et al*., 2012). Briefly, fatty acid and pyruvate oxidation was assessed by measurement of the production of ^14^CO_2_ in a sealed trapping device containing 20 μM palmitate ([1-14C]-palmitate) or 5 mM pyruvate ([2-14C]-pyruvate), tissue sample, and reaction buffer (100 mM sucrose, 10 mM Tris·HCl, 10 mM KPO4, 100 mM KCl, 1 mM 4 MgCl2·6H2O, 1 mM L-carnitine, 0.1 mM malate, 2 mM ATP, 0.05 mM CoA, and 1 mM DTT, pH 7.4) at 37°C.

### Mitochondrial Respiration

Mitochondrial respiration of substrates glutamate and palmitoyl-carnitine was measured in isolated liver mitochondria (n=4, male), as previously described (Morris *et al*., 2013). Briefly, liver mitochondria oxygen consumption was assessed using a Clark-type electrode system (Strathkelvin Instruments, North Lanarkshire, Scotland). Incubations were carried out at 37°C in a 0.5-ml final volume containing 100 mM KCl, 50 mM MOPS, 10 mM K2PO4, 10 mM MgCl2, 0.5 mM EGTA, 20 mM glucose, and 0.2% bovine serum albumin, pH 7.4. Complex I-mediated mitochondrial respiration of either 10 mM glutamate and 10 μM L-palmitoyl-carnitine was monitored in the presences of 1 mM malate and 100 mM ADP. Oxygen consumption (in nmol/min) was normalized to mitochondrial protein in the respirometer cell.

### Indirect Calorimetry Experiments

Energy metabolism was assessed in male and female LPGC1a and WT littermate mice (14-16 weeks of age, n=7-10) for 3 days on either LFD or high-fat/high-sucrose diet (HFHS, D12451: 45% kcal fat, 17% kcal sucrose and 4.73 kcal/g energy density, Research Diets, New Brunswick, NJ, USA) by measuring VO2 in a Promethion continuous metabolic monitoring system (Sable Systems International, Las Vegas, NV, USA), as previously described (Fletcher *et al*., 2018). Animals were acclimated to the indirect calorimetry cages for at least 4 days prior to the start of the dietary intervention and data collection. Animal body weight and food weight were assessed prior to and following the 3-day indirect calorimetry data collection. Rate of energy expenditure (EE) was calculated as EE (kcal/hr) = 4.934 X VO2 (Kaiyala *et al*., 2019), with total EE calculated as the rate of EE times 24 (per day) and summed across the 3-days of analysis. Resting EE was calculated as the average rate of EE (kcal/hr) during the daily 30 minute period with lowest EE times 24 (per day) and summed across the intervention. Non-resting EE is the difference in total- and resting EE. Energy intake was calculated across the 3-day intervention as food intake (grams) times the energy density of the two diets (kcal/g). Energy balance was determined as the difference in energy intake and total EE. As previously described (Abreu-Vieira *et al*., 2015), thermic effect of food was determined as the consensus thermic effect of each macronutrient times the manufacturer provided diet information. From this the thermic effect of food for LFD (D12110704, Research Diets, 3.85 kcal/g, 10% kcals fat, 65% kcal carbohydrate, 20% kcals protein) is 10.5% or 0.4043 kcal/g, and HFHS (D12451, Research Diets, 4.73 kcal/g, 45% kcals fat, 35% kcal carbohydrate, 20% kcals protein) is 8.75% or 0.4139 kcal/g. Activity EE is the difference in non-resting EE and the thermic effect of food. All_meters is an assessment of cage activity and is calculated using the summed distances determined from mouse movements based on XY beam breaks. Cost of movement is a calculation of the energy efficiency of movement and is determined by the activity EE divided by All_meters.

### Common Hepatic Branch Vagotomy (HBV) Experiments

#### Surgery

Starting at 10 – 12 weeks of age, male (n = 7 – 12) and female (n = 6 – 9) LPGC1a and WT littermates underwent common hepatic branch vagotomy or sham surgery under isoflurane as previously reported (Hayes *et al*., 2011). Briefly, a ventral midline incision was made in the abdominal skin and muscle. The ligaments holding the liver lobes were dissected allowing the liver to be everted towards the diaphragm, clearly exposing the esophagus and stomach. The stomach was gently retracted and the common hepatic branch of the vagus was identified as it branches from the descending vagus nerve along the esophagus and runs near the hepatoesophageal artery. For mice in the vagotomy group, the HBV was ablated by cauterization. The abdominal wall and skin were subsequently sutured with 5-0 or 6-0 Vycril. Mice received a single dose of buprenorphine (1.0 mg/kg SQ) for analgesia. Mice were individually housed and daily post-operative monitoring included food intake and body weight assessment for 7 days. Mice had ad lib access to LFD throughout the initial recovery observations and until HFHS weight gain studies began 2 – 3 weeks later.

#### HFHS-induced weight gain and body composition

The one-week HFHS (D12451) feeding studies began 3 – 4 weeks post-surgery. The short-term diet intervention was lengthened from 3-days to one-week as an attempt to decrease variability in outcomes. Body weights, food weights, and body composition was assessed prior to- and post 7 days of exposure to HFHS. Body composition was determined by qMRI (EchoMRI-1100, EchoMRI, Houston, Texas, USA), with fat-free mass being calculated as the differences of body weight and fat mass. As above, energy intake was determined from the food intake and energy density of the diet. Feed efficiency was calculated as the change in body weight divided by the energy intake. Percent metabolic efficiency was calculated as previously described (Luijten *et al*., 2019).

### Statistical Analysis

Data are presented as means and standard error. The two-standard deviation test was used to test for outliers in each group. A two-way ANOVA was used to determine main effects of genotype and diet or genotype and surgery on data separately for each sex. Where significant main effects were observed, post hoc analysis was performed using least significant difference to test for any specific pairwise differences using SPSS version 25 (SPSS Inc., Armonk, New York). Statistical significance was set at p<0.05.

## Results

### Liver-specific PGC1a heterozygous mice have reduced mitochondrial fatty acid oxidation and respiratory capacity

As previously observed (Estall *et al*., 2009), the breeding of female heterozygous floxed *pgc1a* mice with male albumin-cre recombinase mice produced liver-specific, *pgc1a* heterozygous and wild-type (WT) littermate mice (Figure 1A). Female WT mice had >2-fold higher PGC1a mRNA expression compared to males (Figure 1B, p<0.05). LPGC1a male and female mice had 23% and 72% lower liver PGC1a expression compared to WT, respectively (p<0.05). Initial experiments in male mice demonstrate that this reduced liver PGC1a expression led to a ~30% less complete fatty acid oxidation to CO_2_ and ~50% less oxidation of pyruvate to CO_2_ in isolated liver mitochondria (Figure 1C & 1D, respectively, p<0.05) and isolated liver mitochondria (Figure 1D, p<0.05). Additionally, male LPGC1a mice displayed a ~30% reduction in ADP-dependent respiration of glutamate/malate and palmitoyl-carnitine /malate in liver mitochondria compared to WT (Figures 1E & 1F, respectively, p<0.05).

### Reduced liver PGC1a increases susceptibility to short-term HFHS weight gain in male, but not female, mice

To assess whether liver-specific reductions in PGC1a altered energy homeostasis during short-term HFHS exposure, weight gain and energy balance were assessed in male and female LPGC1a and WT littermates following a 3-day HFHS dietary challenge. While females weighed less than males, no difference by genotype was observed in initial body weight within either sex before the 3-day HFHS (data not shown). On the LFD, there was no difference in 3-day weight gain between male WT and LPGC1a mice (Figure 2A). As expected, there was a significant main effect for HFHS-induced weight gain in male mice compared to LFD (p<0.05). Importantly, the male LPGC1a have ~>70% greater weight gain during the 3-days of HFHS exposure compared to WT (p<0.05), resulting in a significant interaction of diet and genotype (p<0.035) (Figure 2A). Interestingly, while female WT mice gained ~1.5-fold more weight during 3-days of LFD compared to LPGC1a (p<0.05, Figure 2B), HFHS feeding resulted in the same weight gain in female mice regardless of genotype (p<0.05).

**Figure 2.**
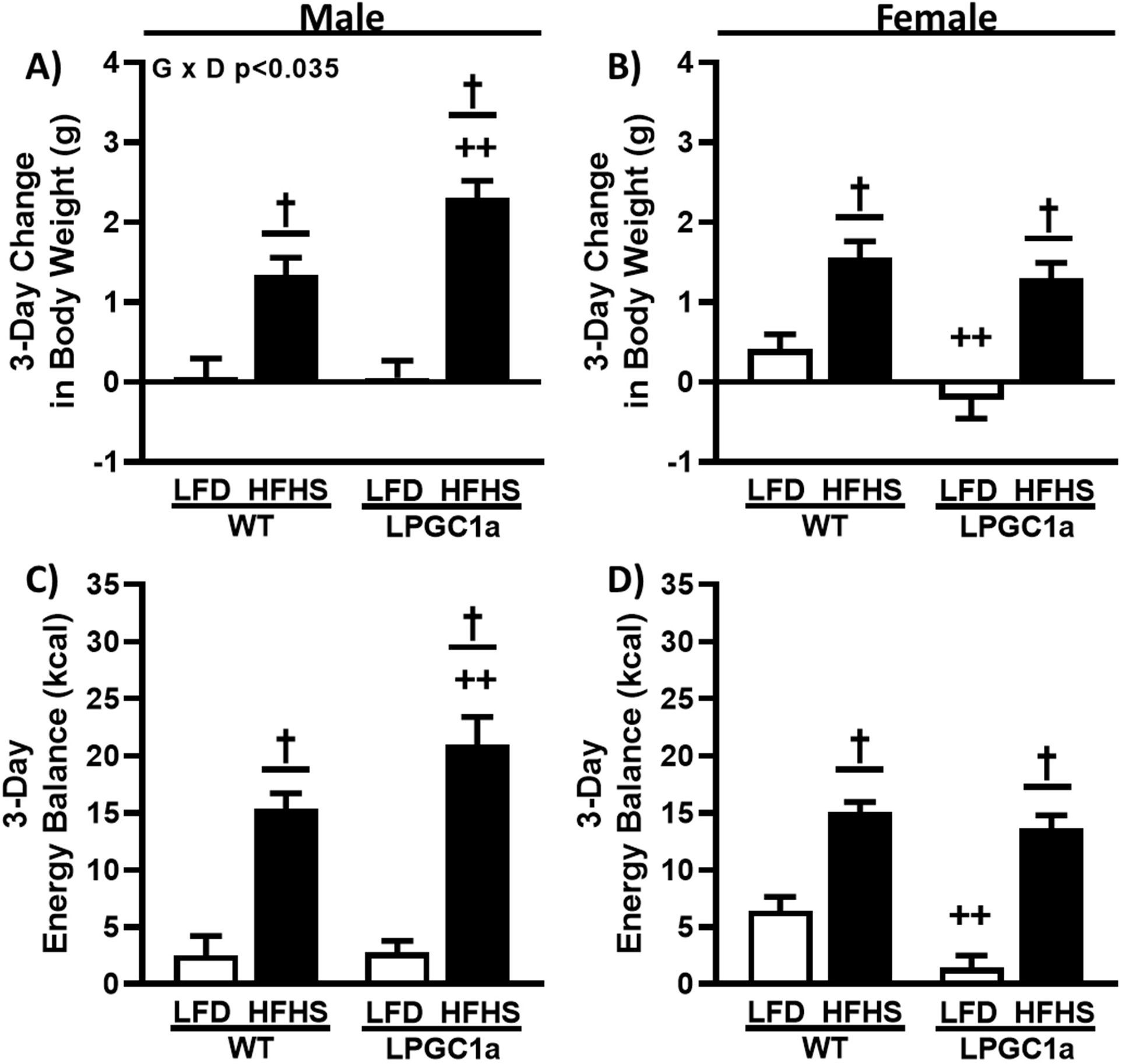
Male LPGC1a mice have greater short-term HFHS-induced weight gain. Body weight gain was assessed during 3-days of LFD or HFHS in male (A) and female (B) WT and LPGC1a mice. Energy balance during the 3-day feeding intervention was determined as energy intake minus total EE for male (C) and female (D) mice. Values are means ± SEM (n=7-10). † p<0.05 main effect of diet, ++ p<0.05 WT vs. LPGC1a within diet.

Energy balance was not different in male mice fed LFD (Figure 2C), with HFHS feeding resulting in a 6- & 8-fold more positive energy balance in WT and LPGC1a male mice, respectively (p<0.05). LPGC1a male mice had a ~35% more positive energy balance during HFHS feeding than WT (p<0.05). As with weight gain, female LPGC1a mice had lower energy balance on LFD compared to WT (~80%, Figure 2D, p<0.05), and shortterm HFHS exposure resulted in more positive energy balance in both female WT and LPGC1a mice (2.5- & 9-fold, respectively, p<0.05). These data demonstrate sex-specific susceptibility to acute diet-induced weight gain in mice with liver-specific reductions in PGC1a is associated with greater impairment in energy homeostasis.

### Male LPGC1a mice have slight increases in HFHS intake and altered feeding behavior

To determine the factors resulting in the greater weight gain and positive energy balance in male LPGC1a mice on HFHS, analysis of energy intake was performed. Energy intake was not different for LFD fed male mice, regardless of genotype (Figure 3A). 3-days of HFHS feeding resulted in ~50% and ~60% greater energy intake in male WT and LPGC1a mice, respectively (p<0.05), with HFHS LPGC1a males tending to have greater energy intake compared to WT (~13%, p=0.09). Female LPGC1a mice tended to have lower energy intake compared to WT (~6%, Figure 3B, p=0.1), and HFHS exposure resulted in a ~40% and ~50% increase in energy intake in female WT and LPGC1a, respectively, compared to LFD (p<0.05). To assess whether any observed differences in energy intake were due to differences in the absolute consumption of the two diets, food intake during the 3-day intervention was analyzed. In Figure 3C, male WT and LPGC1a mice consumed the same amount of LFD, while consuming more HFHS during the short-term exposure (~20% & ~30%, respectively, p<0.05). As expected, based on energy intake data, male LPGC1a tended to have greater intake of the HFHS compared to WT (~13%, p=0.1). Female WT and LPGC1a mice also had increased food intake on the HFHS (~12% & ~20%, respectively, Figure 3D, p<0.05), with LFD LPGC1a mice tending to have reduced intake compared to WT (~10%, p=0.06). These data suggest that the greater weight gain induced by the HFHS in the male LPGC1a mice is partially due to increases in energy intake.

**Figure 3.**
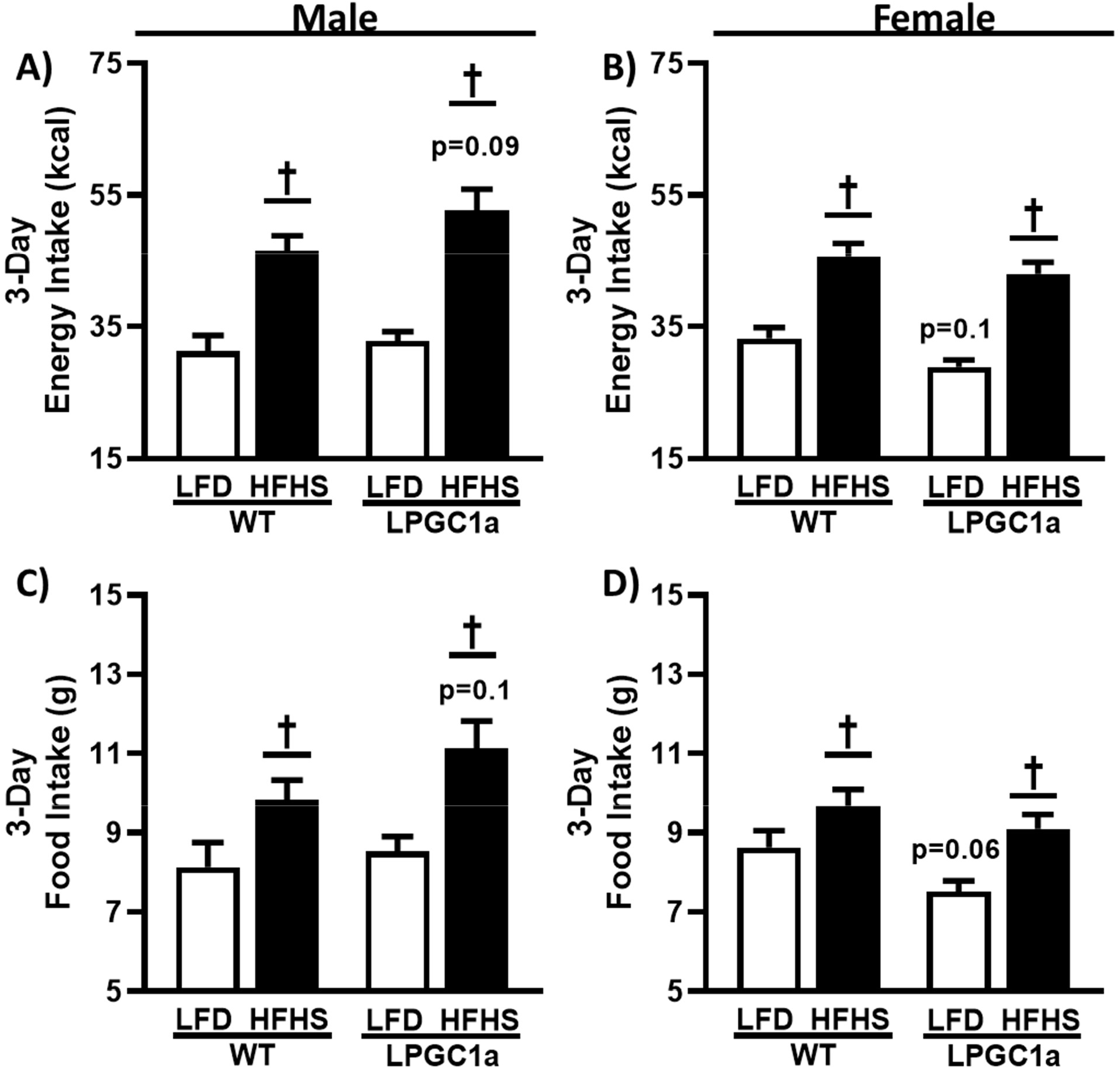
Reduced liver PGC1a subtly impacts HFHS food and energy intake in male mice. A) Energy intake during the 3-days of indirect calorimetry was determined as the energy density of each diet (kcal/g) times the total (B) food intake (g) for male and female WT and LPGC1a mice. Values are means ± SEM (n=7-10). † p<0.05 main effect of diet.

To more specifically investigate the subtle increases observed in food and energy intake only observed between HFHS-fed male LPGC1a and WT mice, we assessed their feeding behaviors during the indirect calorimetry experiments (Figure 4). Male HFHS LPGC1a mice consumed ~35% more food per feeding bout than LFD (Figure 4A, p<0.05), and ~40% more than HFHS WT mice (p<0.05). No difference in grams of food intake per feeding bout were observed between LFD mice. Again, no difference was observed between LFD mice when assessing average number of daily feeding bouts (Figure 4B). However, HFHS WT male mice had a ~40% increase in the number of daily feeding bouts compared to LFD (p<0.05), and a ~30% increase compared to HFHS LPGC1a mice (p<0.05). No difference was observed for the number of feeding bouts between LPGC1a mice on LFD versus HFHS. The length of feeding bouts was not different between LFD fed male WT and LPGC1a mice, however, HFHS feeding resulted in a ~40% reduction in length of feeding bouts for both WT and LPGC1a mice (Figure 4C, p<0.05). Finally, the average time between feeding bouts was reduced ~30% in HFHS male WT mice compared to LFD and HFHS LPGC1a mice (Figure 4D, p<0.05). No difference in length of time between feeding bouts was observed between LFD and HFHS fed LPGC1a male mice. Further, no differences were observed in feeding behavior between female WT and LPGC1a mice (data not shown). Overall, these data demonstrate that reduced liver PGC1a alters feeding behaviors in male mice during short-term exposure to HFHS diet.

**Figure 4.**
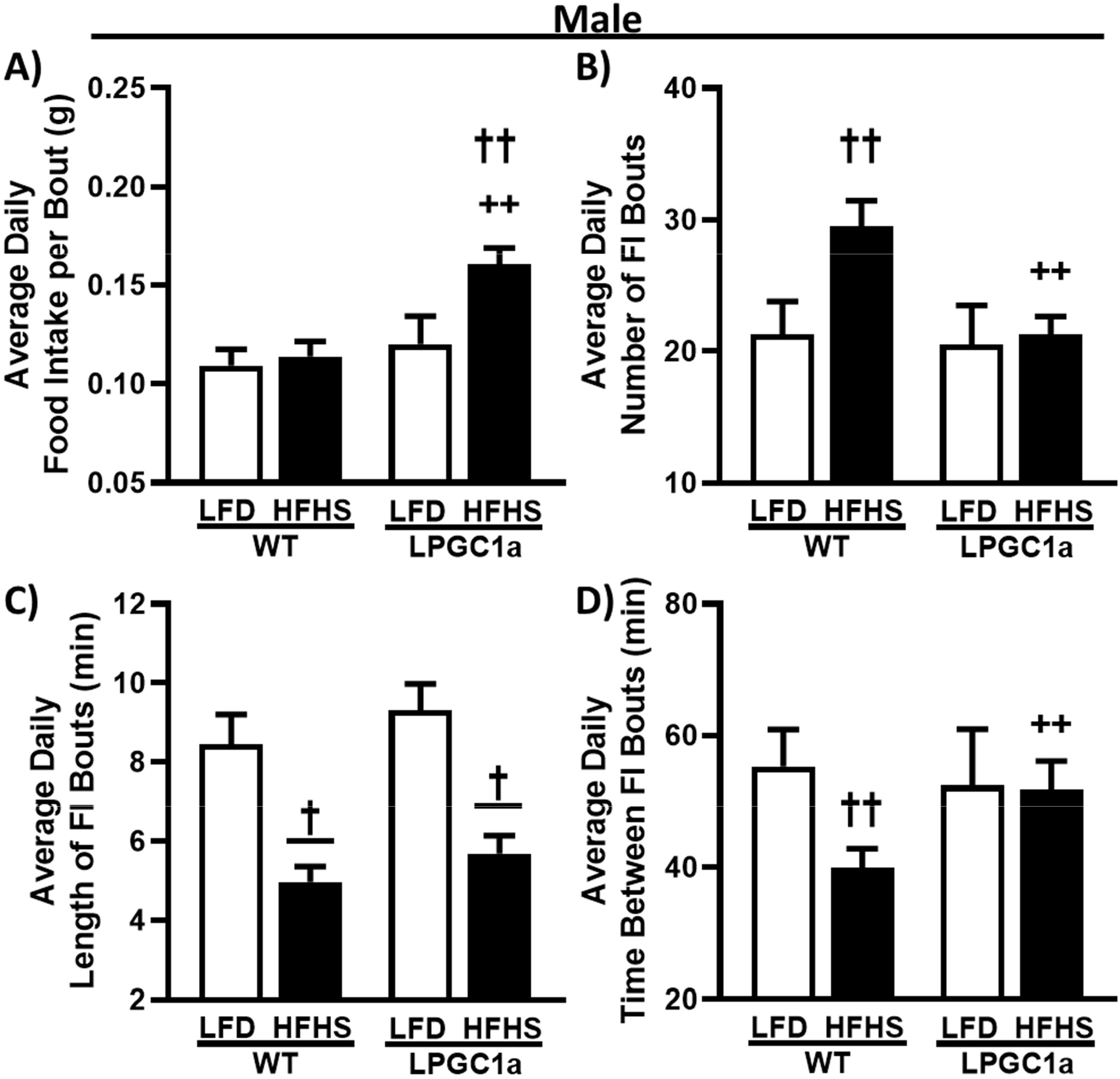
Male LPGC1a mice altered feeding patterns during short-term HFHS feeding. Feeding patterns during the 3-day indirect calorimetry experiments were evaluated in male WT and LPCG1a mice on LFD and HFHS diet for (A) food intake per feeding bout (g), (B) number of daily feeding bouts, (C) length of feeding bouts (min), and (D) time interval between successive feeding bouts (min). Values are means ± SEM (n=7-10). † p<0.05 main effect of diet, †† p<0.05 LFD vs. HFHS within genotype, ++ p<0.05 WT vs. LPGC1a within diet.

### Lower non-resting EE in HFHS-fed LPGC1a male mice prevents HFHS-induced total EE increase

The observed differences in male weight gain in the LPGC1a mice could also be impacted by differences in EE. Therefore, we analyzed total EE and its primary components, resting- and non-resting EE, in male and female WT and LPGC1a mice (Figure 5). Total EE was ~10% greater in male WT HFHS mice compared to LFD (Figure 5A, p<0.05), an effect that did not occur in the LPGC1a mice. Total EE was not different between LFD fed WT and LPGC1a mice regardless of sex (Figure 5A & 5B). Similarly, female WT mice had ~10% greater total EE when fed HFHS compared to LFD (Figure 5B, p<0.05), which was not different from total EE of HFHS LPGC1a female mice. Also, EE was not different between LFD and HFHS in female LPGC1a mice. As the largest component of total EE, resting EE represents the EE of basal metabolic rate and any adaptive thermogenesis. Resting EE was ~10% greater in male WT mice on HFHS compared to LFD (Figure 5C, p<0.05). Male HFHS LPGC1a mice tended to have greater resting EE compared to LFD (p=0.08). Likewise, female HFHS WT mice had ~20% greater resting EE compared to LFD-fed mice (Figure 5D, p<0.05). Again, resting EE tended to be higher in female HFHS LPGC1a mice compared to LFD (p=0.08. Nonresting EE represents 20 - 50% of total EE and is comprised primarily of the thermic effect of food and EE of activity. No difference in non-resting EE was observed between LFD WT males and HFHS, or between LFD males of both genotypes. Interestingly, HFHS feeding reduced non-resting EE in male LPGC1a mice by ~25% compared to LFD (Figure 5E, p<0.05), and tended be lower compared to HFHS WT males (~17%, p=0.08). Together these differences produced a significant interaction (p<0.028) of genotype and diet in male mouse non-resting EE. No difference in non-resting EE was observed in female mice in either genotype or diet (Figure 5F). These data demonstrate that male LPGC1a mice have a mal-adaptive response in non-resting EE upon shortterm exposure to HFHS, which limits or eliminates diet-induced adaptive response of total EE.

**Figure 5.**
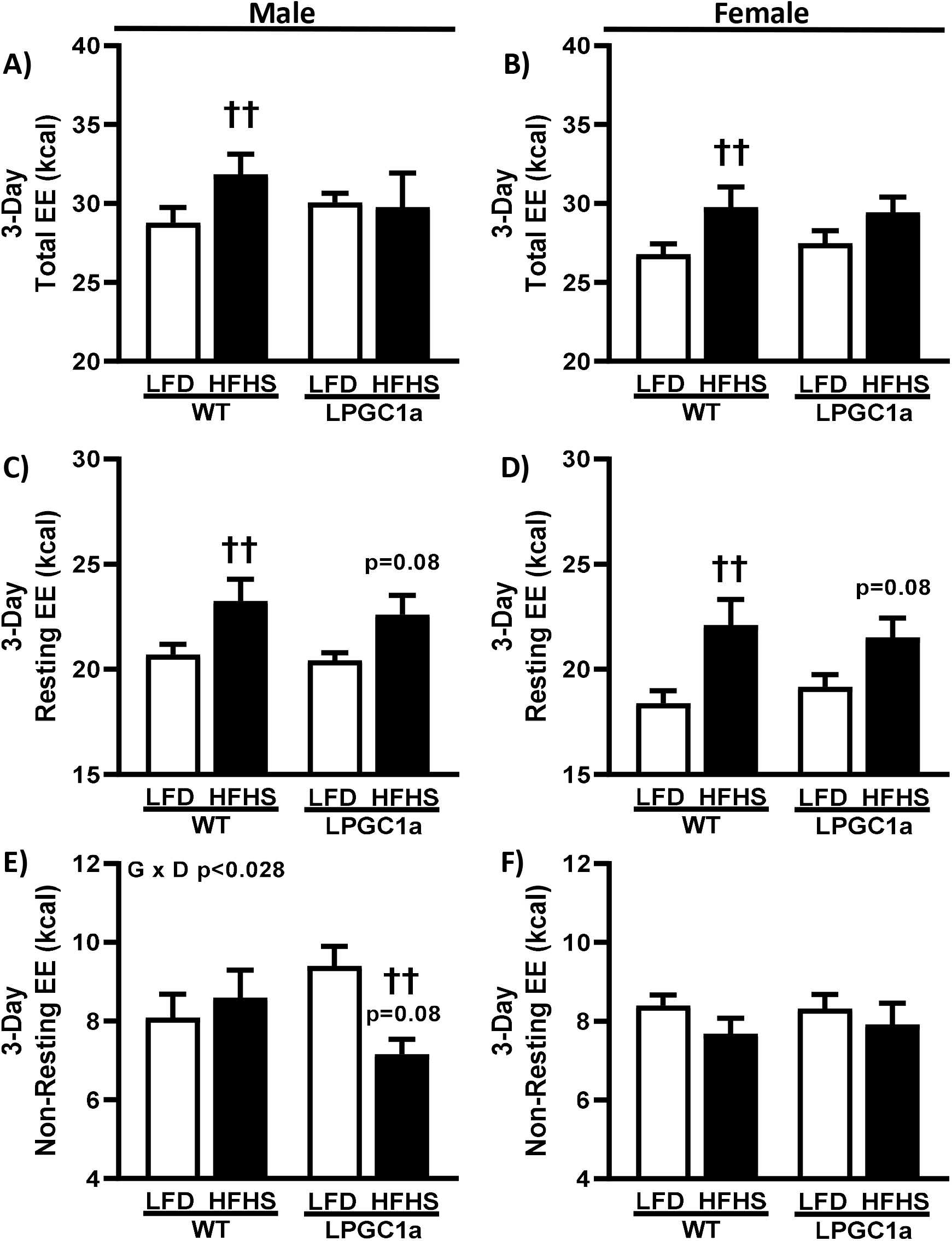
Reduced liver PGC1a blunt HFHS-induced increases in total EE in male mice due to lower non-resting EE. Indirect calorimetry during the 3-day diet interventions was used to determine total EE and it’s primary components: resting EE and non-resting EE in male and female WT and LPCG1a mice. Male and female data is presented for total EE [(A) & (B)], resting EE [(C) & (D)], and non-resting EE [(E) & (F)], respectively. Values are means ± SEM (n=7-10). †† p<0.05 LFD vs. HFHS within genotype.

### Male LPGC1a mice fed HFHS have reduced activity EE and home cage activity

To focus on the diet-induced difference in non-resting EE in male LPGC1a, we also assessed the components of non-resting EE: thermic effect of food and activity EE. The thermic effect of food data mimics food/energy intake (Figure 3), with no differences observed between LFD fed male mice (Figure 6A) and a ~25% and ~35% increase in HFHS fed male WT and LPGC1a mice, respectively (p<0.05). Also, as with food/energy intake, male HFHS LPGC1a mice tended to have greater thermic effect of food compared to WT (p=0.1). Male HFHS LPGC1a mice had ~40% lower activity EE compared to WT and ~50% compared to LFD, which resulted in a significant interaction of genotype and diet (p<0.009) (Figure 6B). Because of differences in activity EE in male HFHS-fed LPGC1a mice we also quantified home cage activity. Male LPGC1a mice on LFD had ~45% greater cage activity compared to WT (p<0.05) (Figure 6C).

**Figure 6.**
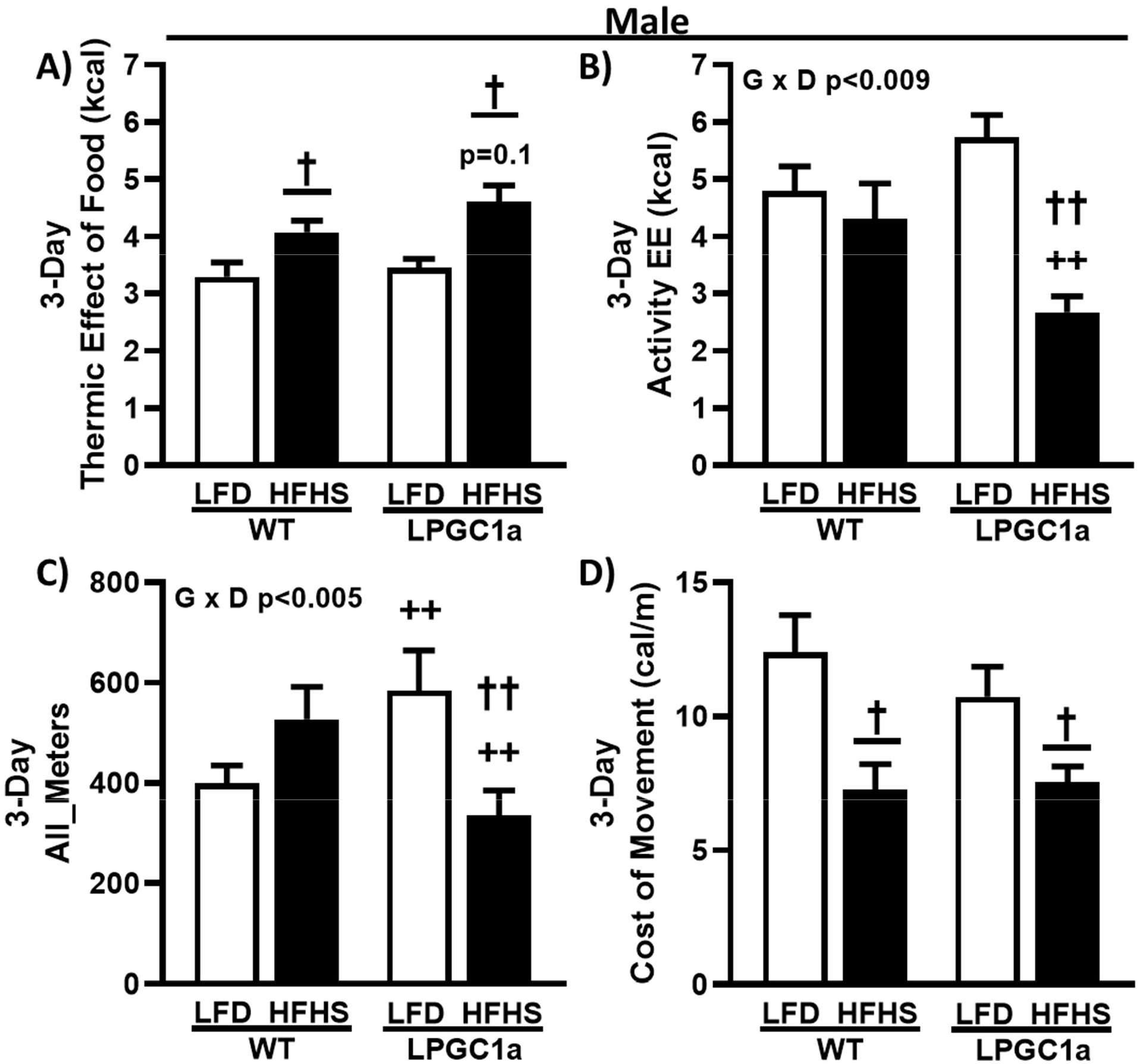
Male LPGC1a mice have lower cage activity and activity EE during short-term HFHS feeding. (A) Thermic effect of food was calculated based on the thermic effect of each macronutrient in each diet multiplied by the amount of food intake. (B) Activity EE was calculated as non-resting EE minus the thermic effect of food. Cage activity is represented as total meters traveled (All_Meters, (C)). The energy efficiency of movement or cost of movement (D) was calculated as the activity EE divided by the cage activity. Values are means ± SEM (n=7-10). † p<0.05 main effect of diet, †† p<0.05 LFD vs. HFHS within genotype, ++ p<0.05 WT vs. LPGC1a within diet.

Importantly, male HFHS LPGC1a mice had ~35% lower home cage activity compared to WT (p<0.05), and ~40% lower compared to LPGC1a LFD (p<0.05). We also calculated energy cost of movement as the activity EE divided by the distance moved during cage activity (Figure 6D). HFHS feeding reduced cost of movement by ~40% and ~30% in male WT and LPGC1a mice, respectively (p<0.05). Finally, no differences were observed in the non-resting EE components between female WT and LPGC1a mice (data not shown). These data demonstrate that reduced liver PGC1a in male mice results in reductions in activity level and associated EE during short-term exposure to HFHS diet.

### Common hepatic branch vagotomy increases short-term HFHS-induced weight gain in wildtype, but not LPGC1a, mice

Previous work has described a hepatic vagal afferent neural pathway that is involved in the regulation of acute food intake (Langhans *et al*., 1985; Horn *et al*., 2001) and insulin secretion (Lee & Miller, 1985; Nagase *et al*., 1993). We performed common hepatic branch vagotomy (HBV) or sham surgery to test if the hepatic vagal afferent pathway is mediating the increased short-term diet-induced weight gain observed in male LPGC1a mice. Following at least two weeks of recovery from the surgical procedure, we assessed body weight gain, energy intake, and changes in body composition for one week (7 days) on LFD followed by 1 week on the HFHS in both WT and LPGC1a male mice. No differences were observed in body weight, energy intake, or body composition between genotype or surgery at the initiation of experiments (data not shown) or during LFD (data not shown). As with the earlier experiments, male sham LPGC1a mice gained ~70% more weight during one-week of HFHS feeding compared to sham WT mice (Figure 7A, p<0.05). Male HBV WT mice gained ~85% more weight during HFHS feeding compared to sham WT (p<0.05), while weight gain in HBV LPGC1a mice was not different from that of sham LPGC1a mice (Figure 7A). Feed efficiency (weight gained divided by energy intake) tracked with the weight gain data (Figure 7B). Feed efficiency in sham LPGC1a mice was ~50% higher than in sham WT mice (p<0.05). HBV WT mice had a ~75% greater feed efficiency compared to sham WT mice (p<0.05). Energy intake during the one-week HFHS was ~12% greater in HBV WT mice and ~9% greater in sham LPGC1a mice compared to sham WT mice (Figure 7C, p<0.05). Body composition was determined in these experiments utilizing qMRI. Short-term HFHS-induced change in fat mass tracked well with energy intake (Figure 7D), with HBV WT mice gaining ~40% fat mass compared to sham WT mice and sham LPGC1a mice tending to have greater fat mass gain than sham WT mice (~30%, p=0.053). Again, HBV in LPGC1a mice did not impact fat mass. Interestingly, short-term HFHS-induced changes in fat-free mass were ~7- and ~2-fold greater in HBV WT and HBV LPGC1a mice, respectively (Figure 7E, p<0.05) compared to sham mice. In addition, there was a ~4.5-fold greater increase in FFM in sham LPGC1a mice compared to sham WT mice (p<0.05). Finally, to more specifically assess the allocation of stored energy, metabolic efficiency was calculated as the energy content of the gained fat- and fat-free mass divided by energy intake (Figure 7F) (Luijten *et al*., 2019). HBV only significantly increased metabolic efficiency in WT mice (~30%, p<0.05), and not in LPGC1a mice. These data highlight that loss of the hepatic vagal afferent neural pathway is sufficient to increase short-term HFHS-induced weight gain in male mice, but does not resolve the male LPGC1a diet-induced weight gain phenotype.

**Figure 7.**
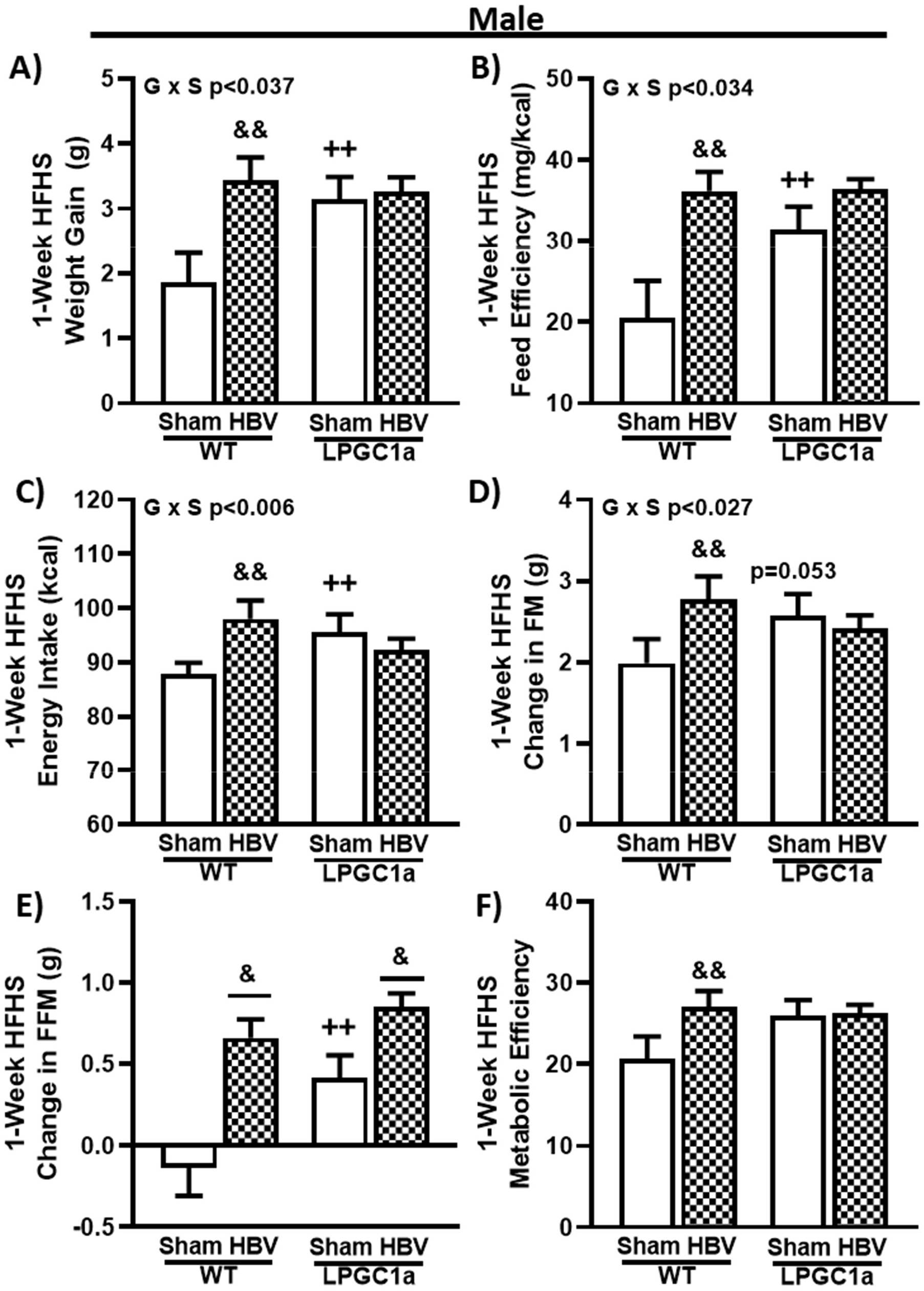
Disruption of liver vagal innervation by common hepatic branch vagotomy increases short-term diet-induced weight gain in wildtype, but not LPGC1a, male mice. Male WT and LPGC1a mice underwent sham or HBV surgery, and were administered a one week HFHS diet following recovery. (A) One week HFHS-induced weight gain. (B) Feed efficiency was determined as the body weight gained divided by the energy intake during the one week HFHS. (C) Energy intake was calculated as the energy density of the HFHS diet multiplied by the one week food intake. Quantitative MRI was utilized to determine fat- and fat-free mass at the initiation and end of the one week HFHS diet, and change in fat mass (D) and fat-free mass (E) are presented. Based on the change in fat- and fat-free mass the (F) metabolic efficiency of weight gain was calculated as the sum of the energy content of fat and fat-free mass multiplied by the respective changes over the one week HFHS, divided by the one week energy intake. Values are means ± SEM (n=7-12). & p<0.05 main effect of HBV, && p<0.05 sham vs. HBV within genotype, ++ p<0.05 WT vs. LPGC1a within surgery.

### Common hepatic branch vagotomy divergently impacts short-term HFHS-induced fat mass gains in female WT and LPGC1a mice

To determine if there are sexually dimorphic contributions of the HBV on LPGC1a-mediated weight gain changes during HFHS feeding, we performed HBV or sham operations in female WT and LPGC1a mice. As with males, there was no difference between HBV and sham at baseline or during LFD. While not significant, HBV produced a divergent HFHS-induced weight gain in female WT and LPGC1a mice (Figure 8A, p=0.088, WT HBV vs. LPGC1a HBV). The observed weight gain was the main effector of the trend toward differences in feed efficiency (Figure 8B, p=0.081, WT HBV vs. LPGC1a HBV), as no difference in one week HFHS energy intake was observed by genotype or surgery (Figure 8C). There was 65% less HFHS-induced fat mass gain in female HBV WT mice compared to sham WT mice (p<0.05), with HBV LPGC1a mice gaining 3.2-fold more fat mass than HBV WT mice (p<0.05). Interestingly, unlike the male mice (Figure 7E), no difference was observed in HFHS-induced fat-free mass gain by genotype or surgery in female mice (Figure 8E). These data suggest that intact hepatic common branch vagal signaling increases macronutrient partitioning to adipose during short-term HFHS feeding in WT female mice, however, loss of this signal interacts with reduced liver PGC1a to increase fat mass gain in female LPGC1a mice.

**Figure 8.**
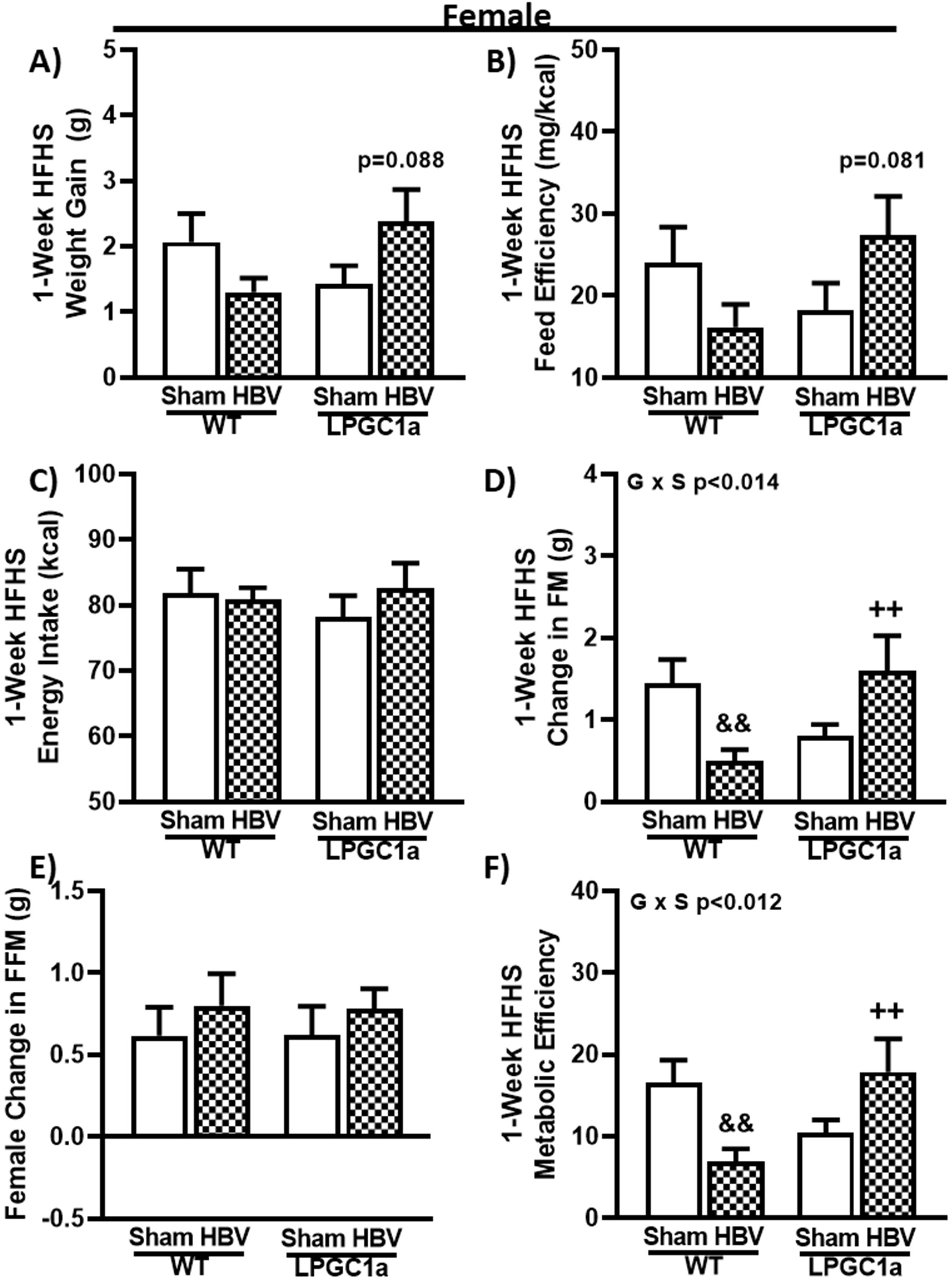
Common hepatic branch vagotomy reduces HFHS-induced fat mass gain in female wildtype, but not LPGC1a, mice. As with males, female WT and LPGC1a mice underwent sham or common hepatic branch vagotomy surgery. (A) Weight gain, (B) feed efficiency, (C) energy intake, change in (D) fat- and (E) fat-free mass, and (F) metabolic efficiency were determined following one week of HFHS feeding. Values are means ± SEM (n=6-9). && p<0.05 sham vs. HBV within genotype, ++ p<0.05 WT vs. LPGC1a within surgery.

## Discussion

Herein, we present novel findings that reduced hepatic PGC1a expression results in increased short-term HFHS-induced weight gain in male, but not female mice. This increased diet-induced weight gain in male LPGC1a mice is associated with altered feeding patterns and reduced activity EE. Further, disruption of hepatic vagal innervation via common hepatic branch vagotomy increased short-term diet-induced weight gain in male WT mice with no observed difference in male LPGC1a mice.

The role of the liver in systemic energy homeostasis has largely been ascribed to the maintenance of blood glucose levels, triglyceride/apolipoprotein metabolism, ketogenesis, and insulin clearance. However, it has previously been observed that liver energy state and vagal innervation are also critical for regulating insulin secretion and acute food intake regulation (32, 45). Further, an inverse relationship between liver-specific modulation of fatty acid oxidation and mitochondrial energy metabolism (Dean *et al*., 2009; Orellana-Gavalda *et al*., 2011; Li *et al*., 2014; Seok *et al*., 2018) or associated secondary changes in liver FAO and mitochondria gene expression has been observed with chronic diet-induced weight gain (Lee *et al*., 2019; Liu *et al*., 2019). Previously, we observed that differences in acute HFHS-induced weight gain, body composition changes, and regulation of food/energy intake were inversely associated with liver tissue PGC1a expression, fatty acid oxidation, and mitochondrial respiratory capacity (Morris *et al*., 2014; Morris *et al*., 2016; Thyfault & Morris, 2017). To more specifically assess the role of liver mitochondrial fatty acid oxidation and respiratory capacity on short-term HFHS-induced changes in energy homeostasis, we utilized a liver-specific, *pgc1a* heterozygous mouse model that has previously been observed to have excess lipid accumulation and reduced expression of fatty acid oxidation and mitochondrial genes (Estall *et al*., 2009).

The mechanistic control of energy homeostasis during each short-term exposure to energy dense foods represents a complex interaction of central and peripheral processes. It has been postulated that liver energy metabolism is involved in this regulation (Friedman, 2007). Herein, we show that liver-specific decreases in fatty acid oxidation and mitochondrial respiratory capacity are associated with increased shortterm diet-induced weight gain in male mice. As expected the 3-day HFHS diet resulted in a more positive energy balance in all groups, which was driven primarily by increased energy intake across the 3-day feeding compared to LFD. However, the increased diet-induced weight gain of the male HFHS-fed LPGC1a mice was supported by a further 35% more positive energy balance compared to WT males. This more positive energy balance results from both a slightly higher energy intake paired with an impaired ability to increase energy expenditure on the HFHS diet. Female LPGC1a mice were not observed to have altered short-term diet-induced weight gain or energy balance compared to WT. Thus, it appears that liver energy metabolism is a peripheral mediator of adaptive homeostatic responses to energy dense diet exposure in male mice.

Previously, we and others have observed reduced food/energy intake in models with increased systemic and liver fatty acid oxidation (Thupari *et al*., 2002; Thupari *et al*., 2004; Morris *et al*., 2014). Earlier findings supported a role for the liver in food intake regulation as an association of chemical inhibition of hepatic fatty acid oxidation and/or reduction in cellular ATP levels with increased acute food intake (Friedman & Sawchenko, 1984; Langhans & Scharrer, 1987; Rawson *et al*., 1994; Friedman *et al*., 1999; Horn *et al*., 2004). However, this work was confounded by potential off-site action of the compounds through intraperitoneal delivery. Herein, our observations of increased HFHS intake and differences in HFHS feeding patterns in a male mouse model of liver-specific alterations in mitochondrial respiratory capacity and fatty acid oxidation suggest the liver is involved in food intake regulation. Interestingly, the reduced number of feeding bouts and increased time between feeding bouts suggests that the male LPGC1a have improved satiety regulation during the HFHS feeding compared to wildtype. While, length of feeding events for both male LPGC1a and wildtype mice is similar, the increased consumption within each feeding bout suggests reduced sensitivity to satiation signals. All told these data implicate liver energy metabolism in the regulation of acute food intake patterns during exposure to HFHS, with further experiments necessary to ascertain why these differences are only observed in males and during HFHS feeding.

Peripheral tissue EE represents the bulk of total EE and has a large capacity to change through alteration in activity EE and adapt to environmental stressors such as cold and diet. The diet-induced adaptive response in EE to energy dense feeding has been proposed as a metabolic response to increased energy intake to minimize positive energy balance excursions (Himms-Hagen, 1984; Lowell & Spiegelman, 2000). This adaptation occurs as increased resting EE through a centrally mediated increase in peripheral non-shivering thermogenesis (Lowell & Spiegelman, 2000). Importantly, both rodent models (Bachman *et al*., 2002; Kazak *et al*., 2017) and humans (Jung *et al*., 1979; Shetty *et al*., 1979) predisposed to diet-induced weight gain and obesity have reduced diet-induced adaptation of EE. As throughout the study female LPGC1a mice have a comparable change in total and resting EE compared to WT during HFHS feeding. However, male LPGC1a mice do not have greater total EE during the HFHS feeding compared to LFD. Interestingly, this is not due to a reduction in diet-induced increase in resting EE, but is due to a substantial reduction in activity EE and cage activity during the 3-day diet intervention. However, it is unclear how acute HFHS feeding reduces activity and activity EE in male LPGC1a mice. While other liver-specific genetic models have demonstrated associated increases in total EE and reduced diet-induced weight gain (Dean *et al*., 2009; Nagata *et al*., 2012), these data represent the first observations of impaired adaptation of total EE due to reduced activity and activity EE during HFHS feeding.

The role of peripheral afferent signaling via the vagus nerve in food intake regulation (reviewed in (Cork, 2018)) and diet-induced obesity (reviewed in (de Lartigue, 2016)) has been well described. Recent data in rats (Banni *et al*., 2012; Yao *et al*., 2018), as well as, in human clinical trials (Shikora *et al*., 2015; Morton *et al*., 2016) implicate vagal stimulation as a treatment for obesity as it reduces body weight, fat mass, and food intake. While food intake regulation is altered through vagal activity by numerous peripheral stimuli, hepatic fatty acid oxidation and energy status has been purported to impact food intake regulation via afferent vagal signals (Langhans & Scharrer, 1987; Horn *et al*., 2001). We performed HBV in LPGC1a mice to assess if vagal signaling of reduced hepatic fatty acid oxidation and mitochondrial respiratory capacity was responsible for the increase in diet-induced weight gain observed. While, the vagotomy had no impact on short-term HFHS-induced weight gain in male LPGC1a mice, it did produce increased weight gain in WT littermates. Previously, HBV produced greater weight gain in rats feed HFD for 3-days compared with sham, with no difference in energy intake per body weight observed (la Fleur *et al*., 2003). More recently, vagal deafferentation resulted in no difference in ad lib low-fat diet food intake and weight gain on chow, but increased food intake and weight gain on HFD (McDougle *et al*., 2020). While the male WT findings support the role of peripheral vagal signaling in diet-induced weight gain, the lack of impact of HBV on male LPGC1a weight gain does not exclude a potential role of hepatic vagal afferents in response to HFHS diets in male LPGC1a mice. Interestingly, HBV in female LPGC1a resulted in greater weight gain compared to WT, which occurred without any change in energy intake. This suggests changes in systemic metabolism and nutrient partitioning in female LPGC1a mice, which is further supported by the observed differences in metabolic efficiency. Further, these findings highlight the need for more sex-specific and comparative studies of the regulation of energy homeostasis via the peripheral vagal neurons. However, these finding also highlight the necessity of more precise deafferentation techniques to more specifically isolate the vagal innervation from the liver to investigate any potential role of liver energy sensing in energy homeostasis regulation.

Limitations. The use of HBV eliminates predominantly peripheral afferent and to a lesser extent, some efferent vagal function for all neurons of the hepatic branch proper and the gastroduodenal branch. This potentially confounds our findings by the loss of afferent gastroduodenal signaling (e.g. – CCK) in food intake regulation and the loss of efferent regulation of hepatic glucose output. Also, the 3-day HFHS studies represent a short exposure time to the diet in mice and may lead to increased variability. Future studies will utilize the one-week HFHS feeding paradigm as utilized in the HBV experiments. Also, the lack of FAO and mitochondrial respiration data in female mice makes it impossible to fully assess any role of these outcomes in the female diet-induced weight gain phenotype. Finally, the study was under powered to allow for direct comparison of males and females. The differences in sex-based response by genotype and HBV to the HFHS exposure highlights the necessity of future studies powered for direct sex comparisons.

In conclusion, we demonstrate that male, but not female, mice with hepatocyte-specific reductions in mitochondrial fatty acid oxidation and respiratory capacity due to heterozygosity of PGC1a are more susceptible to acute diet-induced weight gain during HFHS exposure. This greater weight gain occurred with associated greater energy balance due to a subtle increase in food intake and lack of diet-induced adaptation of total EE. This lack of EE adaptation in male LPGC1a mice resulted from a large drop in activity and activity EE during the HFHS exposure. HBV did not prevent or accentuate this HFHS weight gain in male LPGC1a mice but did result in greater short-term diet-induced weight gain in WT males and LPGC1a females. Overall, these data demonstrate a sex-specific impact of reduced liver energy metabolism on diet-induced weight gain and highlight the need for more specific vagal deafferentation studies and direct sex comparison of interaction of liver energy metabolism and vagal afferent signaling in energy homoeostasis.

## Author Contributions

- E. Matthew Morris was involved in the conception, design, acquisition, analysis, interpretation of the experiments, and the drafting and revision of this manuscript.
- Roberto D. Noland was involved in the design and acquisition of the experiments, and the revision of this manuscript.
- Michael Ponte was involved in the acquisition of the experiments, and the revision of this manuscript.
- Michelle L. Montonye was involved in the acquisition of the experiments, and the revision of this manuscript.
- Julie A. Christianson was involved in the conception and design of the experiments, and the revision of this manuscript.
- John A. Stanford was involved in the conception and design of the experiments, and the revision of this manuscript.
- John M. Miles was involved in the conception and design of the experiments, and the revision of this manuscript.
- Matthew R. Hayes was involved in the conception, design, and interpretation of the experiments, and the revision of this manuscript.
- John P. Thyfault was involved in the conception, design, and interpretation of the experiments, and the revision of this manuscript.

